# TargetGeneReg 2.0: a comprehensive web-atlas for p53, p63, and cell cycle-dependent gene regulation

**DOI:** 10.1101/2021.12.06.470936

**Authors:** Martin Fischer, Konstantin Riege, Robert Schwarz, James A. DeCaprio, Steve Hoffmann

## Abstract

In recent years, our web-atlas at www.TargetGeneReg.org has enabled many researchers to uncover new biological insights and to identify novel regulatory mechanisms that affect p53 and the cell cycle – signaling pathways that are frequently dysregulated in diseases like cancer. Here, we provide a substantial upgrade of the database that comprises an extension to include non-coding genes and the transcription factors ΔNp63 and RFX7. TargetGeneReg 2.0 combines gene expression profiling and transcription factor DNA binding data to determine, for each gene, the response to p53, ΔNp63, and cell cycle signaling. It can be used to dissect common, cell type, and treatment-specific effects, identify the most promising candidates, and validate findings. We demonstrate the increased power and more intuitive layout of the resource using realistic examples.

## Introduction

The cell proliferation cycle and the tumor suppressor p53 are closely linked and represent the most commonly dysregulated signaling pathways in cancer. Despite more than 40 years of research on p53 and many more on the cell cycle, we still lack a comprehensive understanding of the p53 and the cell cycle-dependent regulation of a surprisingly large number of genes. Several mechanisms have been proposed to explain the temporal regulation of hundreds of cell cycle genes (1, 2) and the downstream targets of p53 (3–5), but the substantial overlap between p53 and the cell cycle render the analysis of individual genes difficult.

The expansion of publicly available high-throughput datasets has enabled a more detailed understanding of gene regulatory mechanisms and networks in recent years. We developed a meta-analysis approach to cross-validate results and to improve statistical power by integrating datasets derived from different experimental setups (6). The meta-analysis allows inferring p53 and cell cycle regulation of genes from multiple cell types and treatment conditions and derive common signature genes. It follows the intuitive idea that when multiple datasets agree on a finding it is more likely to be true and that the sum of available evidence provides the best guess for the truth. Previously, we employed our meta-analysis approach to chart the transcriptional programs of the cell cycle, human and mouse p53, the viral oncoprotein E7, and the transcription factor ΔNp63 (6–10). Key findings from our meta-analyses included that p53 serves as a transcriptional activator, while genes repressed by p53 were cell cycle genes at large (6). Specifically, while p53 up-regulates hundreds of genes directly through engaging with chromatin in physical contact with the gene locus, it also down-regulates the large group of cell cycle genes indirectly through its target gene *CDKN1A. CDKN1A* encodes for the cyclin-dependent kinase (CDK) inhibitor p21 that blocks cyclin:CDK complexes leading to the activation of the cell cycle *trans*-repressor complexes DREAM (<u>D</u>P, <u>R</u>B-like, <u>E</u>2F4, <u>a</u>nd <u>M</u>uvB) and RB:E2F (11–14). Moreover, our meta-analyses revealed that the transcription factor complexes RB:E2F, DREAM, and MMB-FOXM1 controlled essentially all of the cell cycle genes, and a small number of genes is both activated by p53 and controlled within the cell cycle by RB:E2F and DREAM (6). However, transcriptome analyses suggest that larger subnetworks of the p53 and cell cycle-dependent gene regulation networks (GRN) are yet to be understood (5, 6).

The target gene regulation (TargetGeneReg) database from our meta-analyses was made available through a web-based atlas at www.TargetGeneReg.org (6) to enable researchers to easily scrutinize the influence of the cell cycle and p53 on any gene of interest. Through www.TargetGeneReg.org, researchers can rapidly determine common as well as treatment, cell type, and species-specific regulations, identify promising targets, and validate findings. It has been used for understanding cell cycle regulators, their signaling cues, and their disease relevance (15–23). Moreover, our database has helped to identify pathways that respond to drugs and stress conditions (24, 25), among many other applications.

Alternative resources such as the p53 BAER hub and the Cyclebase v3.0 either focus on p53-dependent regulation or the influence of the cell cycle on gene expression, respectively (26, 27). However, the integration of both layers of information necessary to understand p53’s contributions to target gene regulation is not readily possible using these tools. Likewise, it is of interest for many researchers to study the target gene expression in other species such as *M. musculus*. While Cyclebase v3.0 includes cell cycle-dependent gene regulation data from other species, the p53 BAER hub does not provide information beyond *H. sapiens*.

The TargetGeneReg resource enabled us to compare the p53 GRN between mouse and human. Surprisingly, up-regulation by p53 displayed substantial evolutionary divergence, while down-regulation of cell cycle genes by p53-p21 is well conserved (9, 28). Moreover, we employed the resource to compare the GRN of p53 to its sibling ΔNp63 and, in contrast to previous reports, we showed that ΔNp63 minimally affects any direct p53 target. Instead, a large number of ΔNp63 targets were cell cycle genes, but the mechanistic link between ΔNp63 and the cell cycle remained unclear (10, 29). Most recently, TargetGeneReg helped us to discover the transcription factor and emerging tumor suppressor RFX7 as a novel node in the p53 GRN, proposing a mechanism for how p53 regulates several targets (30). RFX7 is linked to multiple lymphoid cancers (31), such as Burkitt lymphoma where we and others identified RFX7 as a potential cancer driver (32, 33).

Here, we provide a major update for TargetGeneReg through a substantial expansion of the underlying data resources including more recent RNA-seq and ChIP-seq datasets on p53 and cell cycle regulation, inclusion of data resources on ΔNp63 and RFX7, an upgrade of the website, and visualizations of expanded ChIP-seq data through the UCSC Genome Browser.

## Results

Similar to TargetGeneReg v1.0 (6), TargetGeneReg v2.0 focuses on gene regulation by the tumor suppressor p53 in conjunction with the human cell cycle. An earlier upgrade (TargetGeneReg v1.1) introduced p53-dependent gene regulation and p53 binding data from mouse to TargetGeneReg (9), which, to our knowledge, is unique to the TargetGeneReg resources. Similarly, transcription factor binding data for central transcriptional cell cycle regulators, including the DREAM complex, RB, the MMB complex, and FOXM1, is unique to TargetGeneReg. The upgrade to version 2.0 not only expands the data, but also includes data on p53’s oncogenic sibling ΔNp63 and the emerging tumor suppressor RFX7 (Table 1). In the following sections we provide more detailed information on the data and the applicability of the upgraded TargetGeneReg resource.

**Table 1.** Comparison of TargetGeneReg v2.0 features and resource properties to TargetGeneReg v1.0/1.1 (6, 9), p53 BAER hub (26), and Cyclebase v3.0 (27). Numbers concern data from human except for when ‘mouse’ is indicated. While Cyclebase v3.0 contains multiple datasets on cell cycle-dependent gene regulation from various species, it contains only one dataset from human. ChIP-seq replicates were combined to single datasets for TargetGeneReg but kept separate for the p53 BAER hub. *NA* – not available, information was not provided in the respective publications.

|  | TargetGeneReg<br>v2.0 | TargetGeneReg<br>v1.0/1.1 | p53 BAER<br>hub | Cyclebase<br>v3.0 |
| --- | --- | --- | --- | --- |
| # human genes | **37243** | 18845 | NA | NA |
| human genome version | **hg38** | hg19 | hg19 | NA |
| # p53 expression datasets | **57** | 20 | 16 | - |
| # p53 ChIP-seq tracks/datasets | 32 / 28 | 15 | **41** | - |
| p53RE prediction | **yes** | no | **yes** | - |
| # mouse p53 expression<br>datasets | **15** | **15** | - | - |
| # mouse p53 ChIP-seq datasets | **9** | **9** | - | - |
| # cell cycle expression datasets | **9** | 5 | - | 1 |
| # DREAM ChIP-seq datasets | **17** | 9 | - | - |
| # RB ChIP-seq datasets | **6** | 2 | - | - |
| # MMB-FOXM1 ChIP-seq<br>datasets | **22** | 6 | - | - |
| CHR and E2F motif predictions | **yes** | **yes** | - | - |
| # ΔNp63 expression datasets | **16** | - | - | - |
| # ΔNp63 ChIP-seq datasets | **20** | - | - | - |
| p63RE prediction | **yes** | - | - | - |
| RFX7 target gene prediction | **yes** | - | - | - |
| Genome browser visualizations | **yes** | no | **yes** | no |

### Gene regulation by p53

In the first version of TargetGeneReg, we integrated 20 datasets on p53-dependent gene regulation (6). Since the publication of TargetGeneReg v1.0, several additional high-throughput datasets with varying resolution and experimental strategies became available. To optimally use this additional data and strengthen the power of our resource, we have adjusted our quality control regiment. Specifically, we systematically searched the GEO database for RNA-seq and microarray datasets that employed experimental strategies known to affect p53 signaling. This search included experiments involving MDM2 inhibitors (Nutlin and RG7388), genotoxic and nucleolar stress inducers (Doxorubicin, 5-FU, Actinomycin D, Daunorubicin, Etoposide, Bleomycin, Camptothecin, and UV), viral oncoproteins (SV40 LT, HPV16 E6, and HPV18 E6), exogenous TP53 expression, TNFα, and senescence (oncogene and replication-induced). For inclusion in the updated resource, we required all datasets to comprise at least two biological replicates for both treatment and control conditions. Notably, all datasets we obtained were derived from cell line models. We integrated information from 64 RNA-seq and 35 microarray datasets derived to identify significantly differentially expressed genes. To verify the effects of the selected experiments on known p53-regulated genes, we used a benchmark data set of 116 direct and highly responsive p53 target genes identified earlier based on 16 genome-wide analyses (3). All experiments that yielded less than 50 significantly differentially expressed benchmark targets were removed from further analysis (see Materials and Methods). This measure ensures focus on activation of the p53 pathway by the experimental setup and sufficient power to identify specific p53 activities (Figure 1A). A total of 44 RNA-seq and 13 microarray datasets passed this control (Figure 1B).

**Figure 1.**
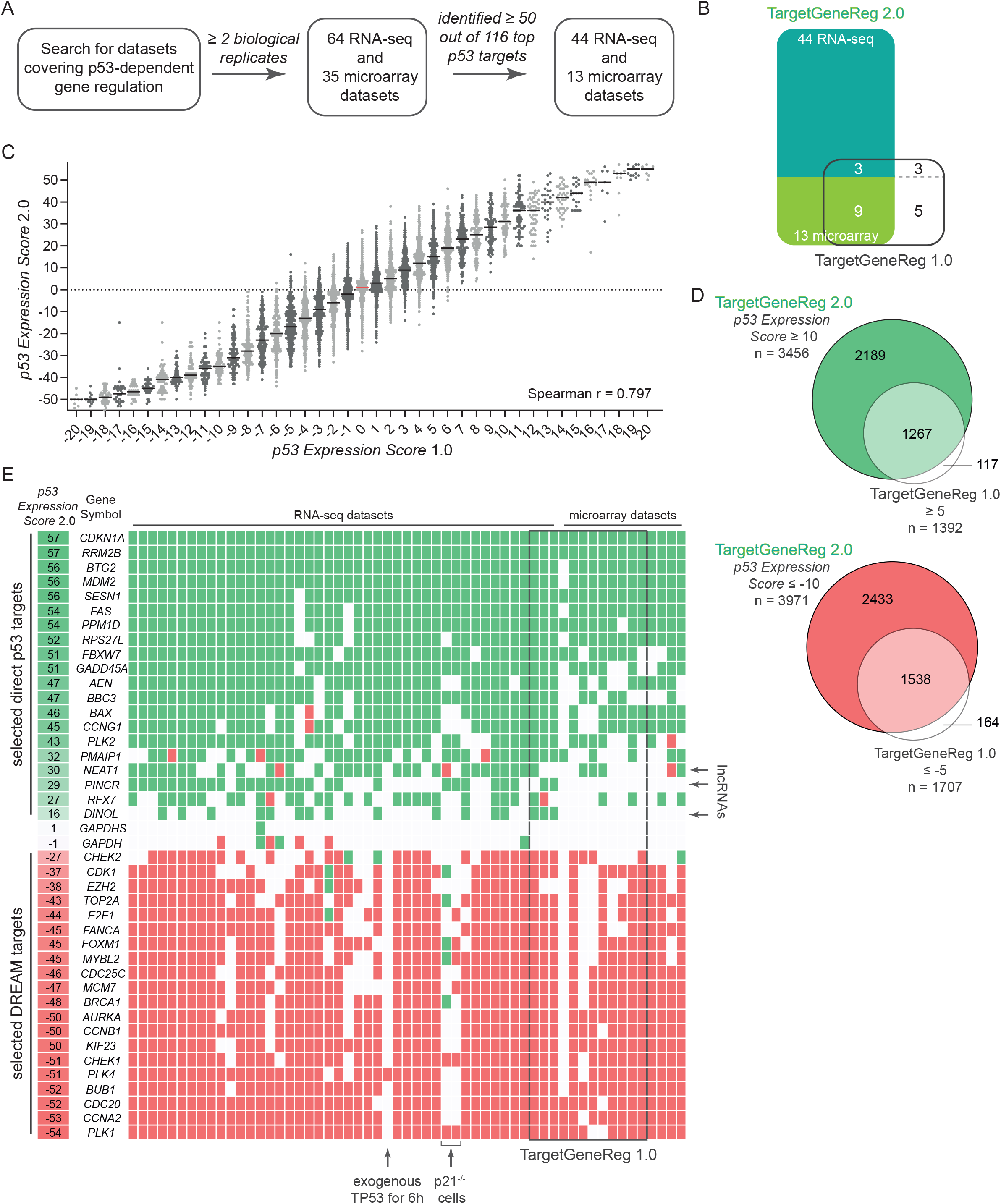
**(A)** Flow chart for the integration of datasets on p53-dependent gene regulation. **(B)** Datasets on p53-dependent gene regulation in TargetGeneReg v2.0 compared to TargetGeneReg v1.0. **(C)** The *p53 Expression Score* v2.0 from TargetGeneReg v2.0 compared to the *p53 Expression Score* v2.0 from TargetGeneReg v1.0 for 17,446 genes present in both databases. Genes are displayed by individual points. The median is indicated by a black line or a red line to highlight ‘0’. **(D)** Genes passing the recommended p53 Expression Score threshold to be considered high-recurrence genes that are up or down-regulated by p53 in TargetGeneReg v2.0 compared to TargetGeneReg v1.0. **(E)** The p53 Expression Score v2.0 and data from the underlying 57 individual datasets visualized for 20 selected direct p53 target genes, 20 selected targets of the DREAM complex, and the non-regulated *GAPDH* genes. It is indicated whether individual datasets were generated using RNA-seq or microarray. Individual datasets that were also present in TargetGeneReg v1.0 are indicated. The lncRNAs *NEAT1, PINCR*, and *DINOL* are highlighted as they were not available in TargetGeneReg v1.0. Details on three datasets in which most DREAM targets are not down-regulated by p53 are shown.

The *p53 Expression Score*, based on these 57 datasets, was calculated for each gene by the number of datasets yielding significant up-regulation minus the number of datasets with significant down-regulation of the gene by p53. To calculate the score, we required a gene to be sufficiently expressed in at least three datasets. A gene was deemed to be expressed when DESeq2 was able to include it in the differential expression analysis, i.e., assign log_2_fold-change and FDR values. A direct comparison of the updated score (*p53 Expression Score* v2.0) with the initial one (*p53 Expression Score* v1.0) exhibits a strong correlation but also suggests an improved resolution (Figure 1C). Most importantly, while the previous version was limited to 18,845 protein-coding genes from hg19, the updated resource now provides a *p53 Expression Score* for 37,243 genes from hg38. The *p53 Expression Score* v1.0 had a minimum threshold of ≥ 5 and ≤ −5 to consider genes with high confidence as being up and down-regulated by p53, respectively (6). In the case of the updated *p53 Expression Score* v2.0, respective thresholds of ≥ 10 and ≤ −10 were passed by 3456 and 3971 genes (Figure 1D). To illustrate the advantage of the updated *p53 Expression Score*, we visualized it together with the underlying individual datasets for 20 selected direct p53 target genes and 20 selected targets of the DREAM complex (Figure 1E).

Of note, some datasets hardly show any down-regulated DREAM targets. This is the case when p21 (CDKN1A) negative cells were used, unable to reactivate the DREAM complex efficiently. Likewise, an experiment where exogenous TP53 was induced for only six hours shows an insufficiency in down-regulating critical cell cycle genes. Moreover, our comparison identifies p53-dependent lncRNAs such as *DINOL* (34), *PINCR* (35), and *NEAT1* (36, 37) excluded from the previous version because of insufficient data (Figure 1E). In addition to information on the p53-dependent regulation of thousands of non-coding RNAs, the updated *p53 Expression Score* v2.0 provides much more detailed information on p53-dependent regulation for hundreds of genes for which the *p53 Expression Score* v1.0 was inconclusive. Consequently, 1674 and 1917 genes that previously displayed a *p53 Expression Score* v1.0 between 5 and −5 now passed the threshold of a *p53 Expression Score* v2.0 of ≥ 10 and ≤ −10, respectively, indicating a differential regulation by p53 with high recurrence.

The direct binding of p53 to the gene promoter is a crucial property of many genes up-regulated by this transcription factor (3–5). While TargetGeneReg v1.0 integrated 15 datasets on p53 genome binding (6), the updated version now integrates an expanded collection of 28 ChIP-seq datasets. While the previous version just displayed the number of datasets that identified p53 binding near a gene’s TSS, the updated resource contains precise binding location information and visualizations thereof. To enable users to rapidly visualize the large number of 28 individual p53 ChIP-seq datasets, we provide a ‘peak-of-peaks’ data track representing a pile-up of p53 peak regions from individual datasets (Figure 2A). Therefore, the ‘peak-of-peaks’ track provides quick summary information on how many datasets identified p53 binding to any locus in the genome. In addition, the p53 response element (p53RE) most closely resembling a canonical p53RE is displayed for each peak-of-peaks, as described previously (10). In addition, the ChIP-seq tracks from all individual datasets can be displayed upon selection, providing a seamless visualization of cell type and treatment-specific information next to the summary data (Figure 2A and B). As established previously, any binding site with support from at least five datasets is considered to be of high recurrence (9, 10). The website’s ‘Overview’ section indicates for every gene whether it displays a high-recurrence binding site within 2.5 kb of a TSS, and whether a high-recurrence binding site is linked to the gene locus through a double-elite enhancer:gene association listed in the GeneHancer database (38).

**Figure 2.**
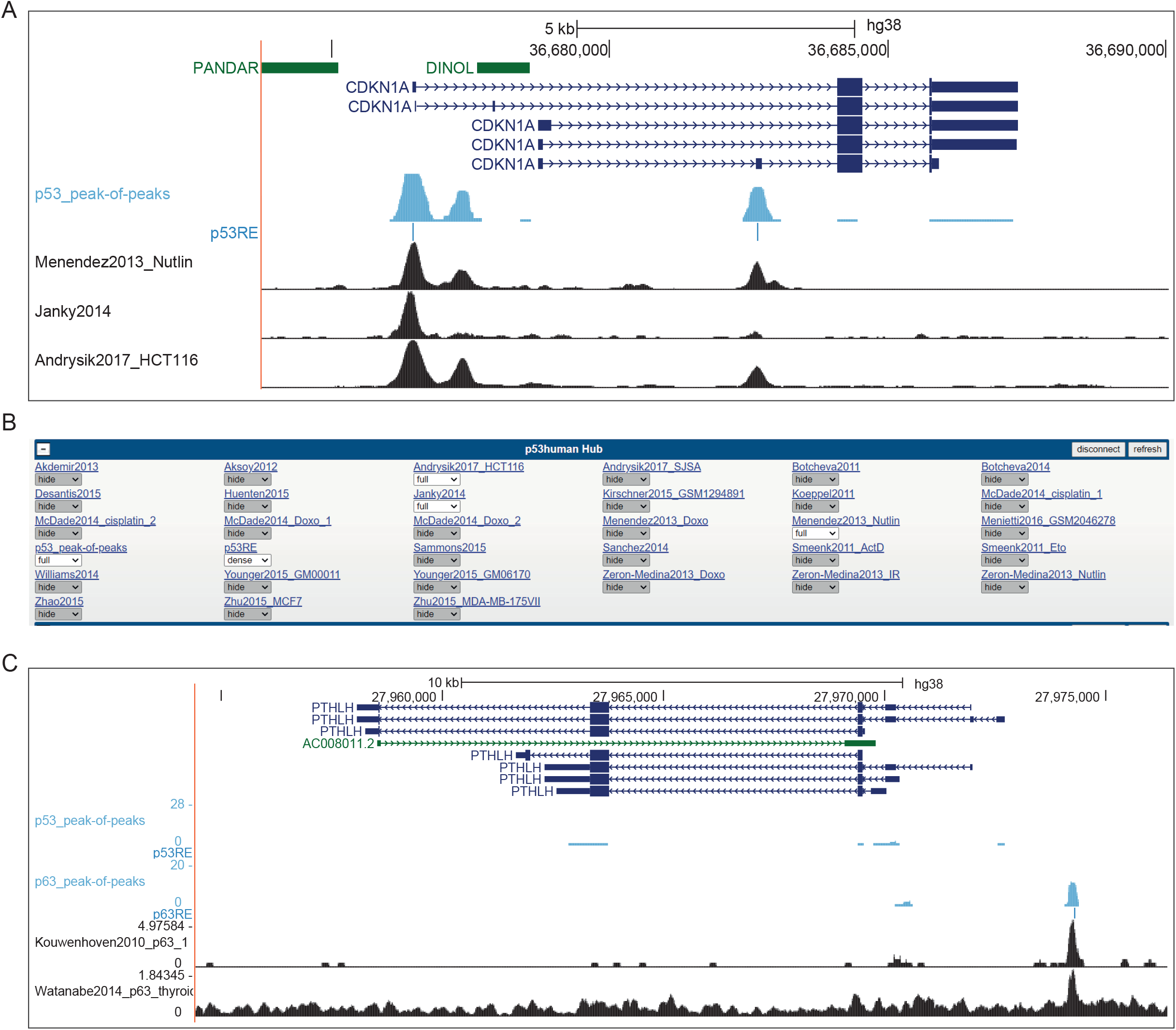
**(A)** Image of UCSC Genome Browser displaying the *CDKN1A* locus linked from TargetGeneReg v2.0. The blue tracks display the p53_peak-of-peaks summary and the most likely underlying p53RE that has been identified. Individual p53 ChIP-seq tracks can be selected from **(B)** the track hub that is loaded through the TargetGeneReg v2.0 linkage. Together, the visualization provides precise visualization of p53 binding sites locations and their underlying p53RE and enables seamless comparisons between summary data and individual datasets. **(C)** Image of UCSC Genome Browser displaying the *PTHLH* locus linked from TargetGeneReg v2.0. The blue tracks display the p53_peak-of-peaks and p63_peak-of-peaks summaries and the most likely underlying p53RE and p63RE. Individual p63 ChIP-seq tracks can be selected from the track hub hat is loaded through the TargetGeneReg v2.0 linkage, as shown for p53 above. *PTHLH* is a direct target of ΔNp63 but not p53, and the unique p63 binding site can be readily seen by comparing the peak-of-peaks summary binding data.

Together, TargetGeneReg v2.0 provides information on p53-dependent gene regulation for twice as many genes from almost three times as many datasets in total and almost five times as many datasets that follow the tightened control measures. In addition, it provides almost twice as many p53 ChIP-seq datasets, predictions for the underlying p53RE, and precise location visualizations.

### Cell cycle-dependent gene regulation

Cell cycle genes play essential roles in cell cycle progression and therefore are typical markers of proliferation that are dysregulated in most cancers (39). Like many other growth restricting processes, the tumor suppressor p53 down-regulates cell cycle genes to sustain cell cycle arrest. Based on TargetGeneReg v1.0, we consolidated the five cell cycle gene peak clusters defined by Whitfield *et al*. (40) to two major groups of cell cycle genes, namely G1/S and G2/M genes (6). Here, we expanded the previous resource’s five datasets to include four additional datasets (Figure 3A). For all genes identified as cell cycle-dependently regulated in at least three of the nine datasets, we predicted whether the gene is a G1/S or a G2/M gene based on a majority vote by the nine datasets (see Materials and Methods). The website’s ‘Overview’ section provides information on the number of datasets that suggest a gene to be driven by the cell cycle (‘Cell Cycle Expression Score’) and its classification prediction (‘Cell Cycle Gene Category’).

**Figure 3.**
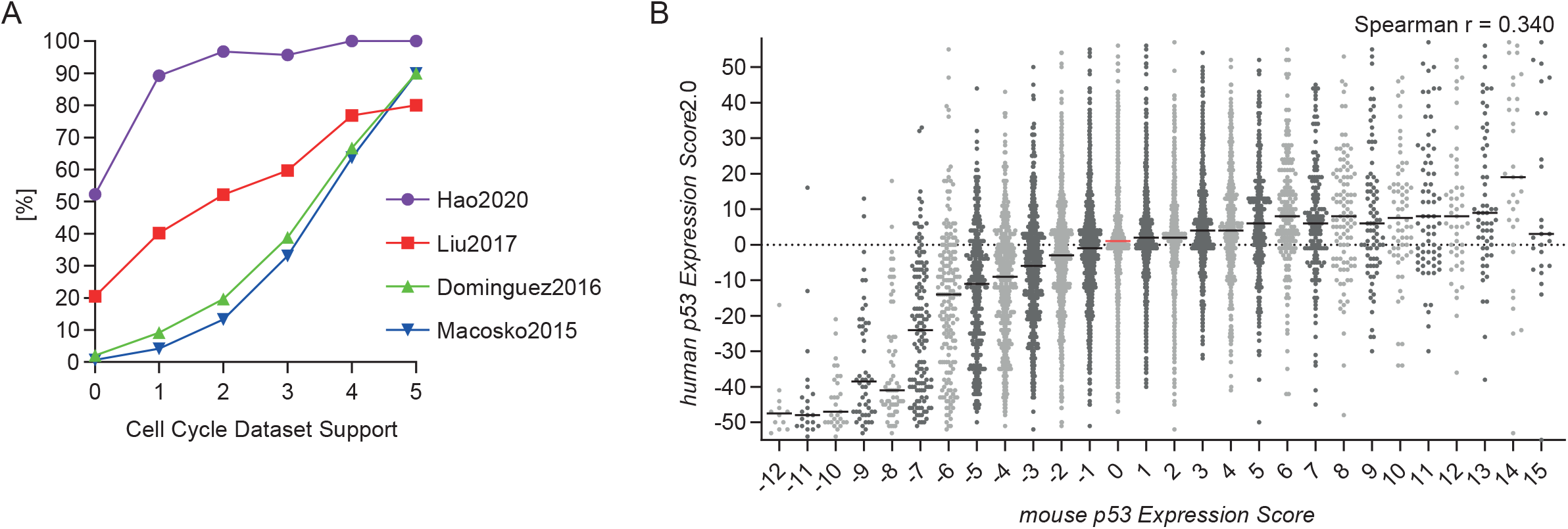
**(A)** Four new datasets on cell cycle-dependent gene expression have been added (43–46). Following our previous quality control (6), we tested whether the datasets were more likely to identify a gene as cell cycle gene when more datasets from TargetGeneReg v1.0 (X-axis) agreed on its cell cycle gene status. **(B)** The *mouse p53 Expression Score* (9) compared to the *human p53 Expression Score* v2.0 for 14,712 one-to-one orthologs with both scores. Genes are displayed by individual points. The median is indicated by a black line or a red line to highlight ‘0’.

The two distinct groups of G1/S and G2/M genes are primarily characterized by E2F and CHR (<u>c</u>ell cycle genes <u>h</u>omology region) DNA recognition motifs in their promoters, respectively (2, 6, 41). The DREAM complex can bind to both E2F and CHR motifs through its respective subunits E2F4 and LIN54, while RB:E2F specifically binds G1/S cell cycle genes through E2F motifs. Alternatively, the transcription factors B-MYB (also known as MYBL2) and FOXM1 associate with DREAM’s LIN54-containing MuvB core complex later in the cell cycle to specifically drive the expression of G2/M genes through binding their CHR sequences (2). To allow a more comprehensive analysis of cell cycle-dependent regulation, we expanded the nine datasets on genome binding by DREAM complex components to 17. Similarly, we extended the two previous datasets on RB binding to six datasets, and the previous six datasets on MMB:FOXM1 (B-<u>M</u>YB:<u>M</u>uv<u>B</u>:FOXM1) binding to 22 datasets. Potential E2F and CHR motifs under respective RB and MMB:FOXM1 binding sites have been predicted using HOMER (42). The individual ChIP-seq tracks, peak-of-peaks, and motif predictions are available through UCSC Genome Browser visualizations.

### Gene regulation by mouse p53 and its difference to human p53

Previously, we employed our meta-analysis approach on mouse p53 synthesizing p53-dependent gene regulation data across 15 datasets, and we made the data available through TargetGeneReg v1.1 (9). Here, we integrated our database on p53-dependent gene regulation in mice (mm10) with the updated database on p53-dependent gene regulation in humans (hg38) described above. Therefore, TargetGeneReg v2.0 includes mouse p53-dependent gene regulation data for the one-to-one orthologs of 14,712 human genes. While there is a good correlation between the *mouse p53 Expression Score* and the *human p53 Expression Score* v2.0 for genes down-regulated by p53, the correlation for up-regulated genes is poor (Figure 3B), indicating a strong and a comparably low evolutionary conservation of p53 down and up-regulated genes, respectively. Similar results have already been reported for the first *p53 Expression Score* (9, 28). Precise binding data (ChIP-seq) of mouse p53 is available through links to the UCSC Genome Browser with embedded track hubs similar to human protein binding data described above.

### Gene regulation by ΔNp63

We previously employed our meta-analysis approach to provide a comprehensive resource for gene regulation by p53’s sibling ΔNp63, an essential oncoprotein in squamous cell carcinomas (10). Given its relevance to cancer and close connection to p53, we integrated our ΔNp63 database comprising 16 datasets on p63-dependent gene regulation and 20 ChIP-seq datasets on p63 DNA binding into TargetGeneReg v2.0. The Δ*Np63 Expression Score* and predictions of p63 targets (180 high-recurrence targets available in Table 1 from Riege *et al*. (10)) and potential p63 targets (comprising all genes bound and regulated by p63) are available through the website’s ‘Overview’ section. DNA binding data and identified p63 response elements (p63RE) are available through UCSC Genome Browser track hubs and enable a seamless comparison between individual p63 ChIP-seq tracks, summary data thereof (p63_peak-of-peaks), and respective data from p53. *PTHLH*, for instance, is a direct target gene of ΔNp63 but not of p53 (Figure 2C).

### Expanding the p53 gene regulatory network through RFX7

Complex cross-talk between signaling pathways impedes the identification of indirect gene regulatory mechanisms employed by p53. For example, following two decades of conflicting data on mechanisms of p53-dependent gene repression p53 was found to serve as a transcriptional activator that represses genes indirectly, with its target p21 taking a predominant role through its profound influence on down-regulating the cell cycle genes (3, 4, 6, 7). Recently, we identified the transcription factor and emerging tumor suppressor RFX7 as a vital node in the p53 transcriptional program (30). RFX7 orchestrates a subnetwork of tumor suppressor genes in response to cellular stress and p53. Given the crucial role of the novel p53-RFX7 signaling axis in the p53 gene regulatory network and its potential importance to cancer biology, we included the data on RFX7 target genes in TargetGeneReg 2.0. The website’s ‘Overview’ section displays whether a gene has been predicted as an RFX7 target, offering a mechanistic explanation for its p53-dependent up-regulation. In addition, RFX7 ChIP-seq tracks are available through the UCSC Genome Browser visualizations.

### Navigating the TargetGeneReg 2.0 website

Our updated resource is tailored to rapidly provide information on genes of interest entered in the main input field. The arguably strongest asset of the TargetGeneReg resource is summary data on gene regulation by various transcription factors generated through a synthesis of multiple individual datasets integrated by our meta-analysis approach. Therefore, the ‘Overview’ section situated at the top of the one-page website provides all summary information on the genes of interest that have been entered (Figure 4). Importantly, it enables a quick direct comparison of the summary data between multiple genes. More detailed information on gene regulation from the individual datasets is provided in the detailed sections below, which are equipped with sorting options to quickly identify the most relevant data. Precise transcription factor binding data visualized through the UCSC Genome Browser (47) are available through the genomic position links provided in the ‘Overview’ section for both human (hg38) and mouse (mm10).

**Figure 4.**
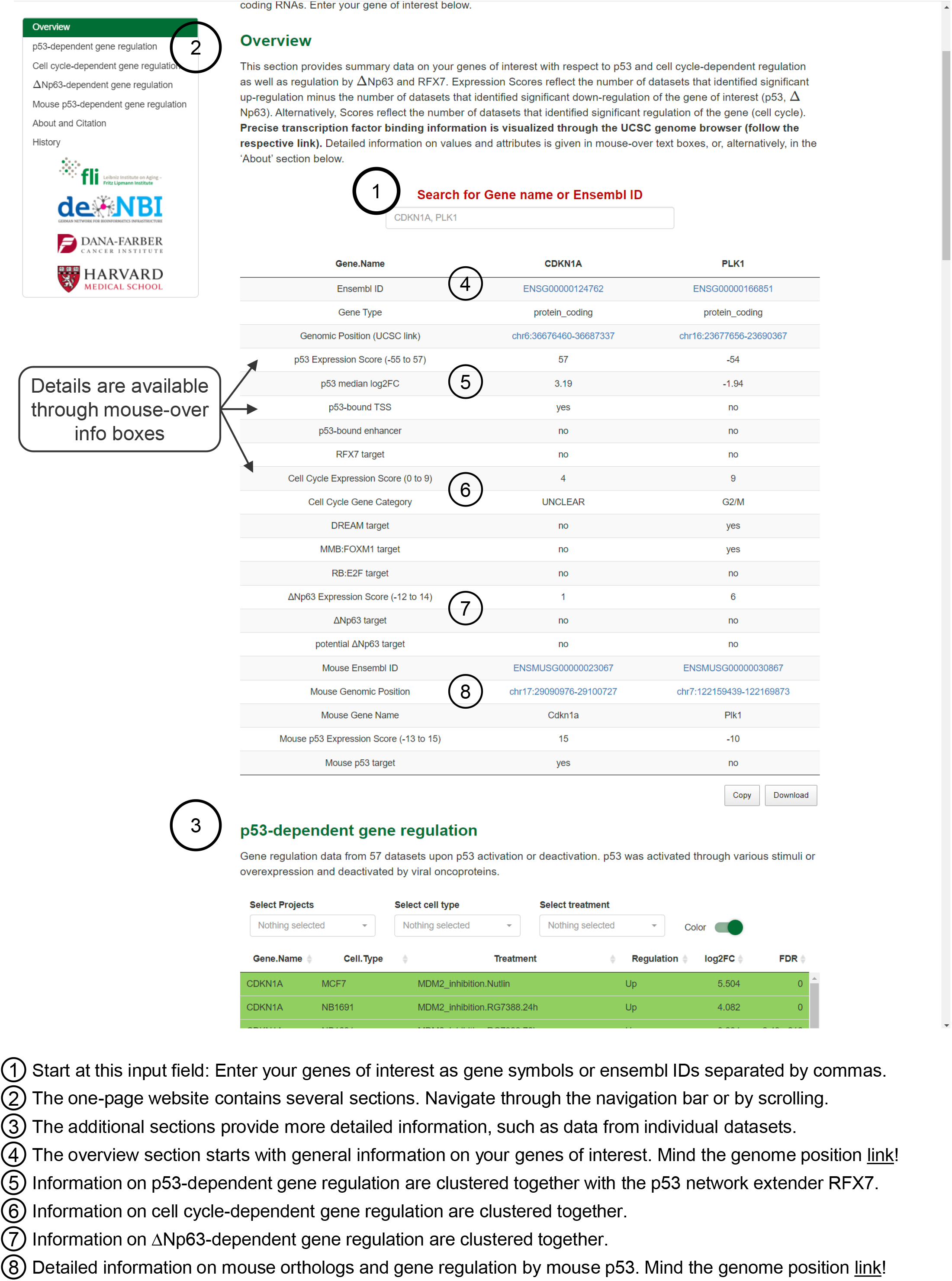
The TargetGeneReg 2.0 website design. Start by entering your genes of interest as gene symbol or Ensembl gene ID. The overview section provides a helpful summary on the regulation of your genes of interest. Details on each point in the overview section is available through mouse-over boxes and through the ‘About’ section. The additional sections are available through the navigation bar in the upper left corner or by scrolling through this one-page website design. The additional sections contain more detailed information, such as data from the individual datasets and citation and history data for the resource.

## Discussion

The TargetGeneReg has gained a strong reputation in the p53 and cell cycle communities. It provides a deeper insight into p53 and cell cycle-dependent gene regulation mechanisms. The presented upgrade, TargetGeneReg v2.0, substantially improves this resource. Like its predecessor, the resource is tailored to quickly retrieve information on the users’ genes of interest and provides swift comparisons between genes and experimental conditions. TargetGeneReg v1.0 was a starting point to help the p53 and cell cycle communities to gain deep biological insights by providing reference points and a platform that enables users to quickly test whether their genes of interest are likely regulated by p53 or the cell cycle. The new version integrates more datasets, substantially improving its power. Specifically, TargetGeneReg now includes data on non-coding RNAs and provides information on the gene regulation by p53’s oncogenic sibling ΔNp63 and the emerging tumor suppressor RFX7.

Visualization of the transcription factor binding data through the UCSC Genome Browser provides precise location information for the user to better interpret the potential consequences of the binding for their gene of interest. For example, p53 binding to intronic locations can induce alternative transcription start sites leading to transcript variants with shortened 5’ sequences, as reported for *MDM2* and *FBXW7* (48, 49). Visualization of the strongest scoring underlying p53RE and p63RE provides an unprecedented depth of binding information.

While the summary data, such as the *Expression Scores* are particularly helpful to quickly assess the regulation of genes, it is critical to tally the characteristics of individual datasets and genes used for the generation of this summary. Importantly, a low *Expression Score* does not rule out p53 or cell cycle-dependent regulation. In addition to biological variability, such as cell line-specific differences, genes may evade the differential expression detection due to low transcript abundances, low but biologically relevant fold-changes, or methodological limitations (e.g., limited number of replicates and sequencing depth). For example, many non-coding RNAs have low expression levels and thus may have a low *p53 Expression Score* although they are actually strongly regulated by p53 (e.g., *DINOL* displayed in Figure 1E). From a statistical perspective, this situation would require the integration of more data sets to increase the statistical power. To address this limitation in spite of additional datasets, we display the ‘p53 median log2FC’. The combination of a low *p53 Expression Score* and a high ‘p53 median log2FC’ might indicate that a gene evades differential expression detection due to a low overall expression level.

Together, with TargetGeneReg 2.0 we provide a comprehensive resource on p53-dependent regulation in humans and mice. Additional information on ΔNp63 and cell cycle-dependent gene regulation facilitates the discovery of further novel biological insights.

## Materials and Methods

### RNA-seq analysis pipeline

We used Trimmomatic (50) v0.39 (5nt sliding window approach, 5’ leading and mean quality cutoff 20) for read quality trimming according to inspections made from FastQC (https://www.bioinformatics.babraham.ac.uk/projects/fastqc/) v0.11.9 reports. Illumina adapters as well as mono-and di-nucleotide content were clipped using Cutadapt v2.10 (51). Potential sequencing errors were detected and corrected using Rcorrector v1.0.4 (52). Ribosomal RNA (rRNA) transcripts were artificially depleted by read alignment against rRNA databases through SortMeRNA v2.1 (53). The preprocessed data was aligned to the reference genome hg38, retrieved along with its gene annotation from Ensembl v102 (54), using the mapping software segemehl (55, 56) v0.3.4 with adjusted accuracy (95%) and split-read option enabled. Mappings were filtered by Samtools v1.12 (57) for uniqueness and, in case of paired-end data, properly aligned mate pairs. Differential gene expression and its statistical significance was identified using DESeq2 v1.30.0 (58). Common thresholds of |log_2_fold-change| ≥ 0.25 and FDR < 0.05 were used to identify significantly differentially expressed genes.

### Microarray analysis pipeline

All microarray datasets were available at a pre-processed stage at the Gene Expression Omnibus (GEO) and we re-analyzed the datasets with GEO2R to obtain fold expression changes and Benjamini Hochberg-corrected p-values (FDR) (59). Gene identifiers were mapped to Ensembl Gene IDs using the Ensembl annotation data v102 (54). Common thresholds of |log2fold-change| ≥ 0.25 and FDR < 0.05 were used to identify significantly differentially expressed genes.

### Meta-analysis / generation of Expression Scores

Following our meta-analysis approach (6), *Expression Scores* for genes regulated by human and mouse p53 and ΔNp63 were calculated as the number of datasets that find the gene to be significantly up-regulated minus the number of datasets that find the gene to be significantly down-regulated by the respective transcription factor. Both, the *mouse p53* and the Δ*Np63 Expression Score* were published previously (9, 10). The *Cell Cycle Expression Score* reflects the number of datasets that identified a gene as cell cycle-regulated gene. The *Cell Cycle Gene Category* is calculated by a majority vote of the nine datasets on cell cycle-dependent gene expression and is displayed for each gene that shows a *Cell Cycle Expression Score* ≥ 3. Precisely, each dataset that identified peak expression of the gene in ‘G1’, ‘G1/S’, or ‘S-phase’ is grouped as ‘G1/S’, and peak expression in ‘G2’, ‘G2/M’, ‘M’, and ‘M/G1’ is grouped as ‘G2/M’.

### ChIP-seq data integration

Peak datasets and bigwigs from ChIP-seq experiments were retrieved from CistromeDB (60) ensuring a common data processing pipeline and thereby a direct comparability. Only RFX7 ChIP-seq data were taken from our recent study (30) as they were not yet available through CistromeDB. Bigwigs (ChIP-seq tracks) have been made available through track hubs for the UCSC Genome Browser (47). Notably, while ChIP-seq replicates are available as individual tracks in the track hubs, they have been jointly considered as one dataset for the generation of peak-of-peaks summaries. Precisely, when replicate experiments were available, all peaks were used that have been identified in at least two replicates. To identify overlapping and non-overlapping peaks Bedtools ‘intersect’ was employed, and to generate the peak-of-peaks summaries, multiple peak files were combined using Bedtools ‘multiinter’ (61). The p53 and p63 ChIP-seq collections and summaries on human and mouse p53 and ΔNp63 have been published previously (9, 10). Similarly, p53REs and p63REs were taken from our previous study (10).

## Acknowledgements

This work was supported by the German Federal Ministry for Education and Research (BMBF) [031L016D to S.H.] and in part by grants from the National Institutes of Health [R35CA232128 and P01CA203655 to J.A.D.]. The FLI is a member of the Leibniz Association and is financially supported by the Federal Government of Germany and the State of Thuringia.

We gratefully acknowledge Bernd Senf from the FLI Core Facility Life Science Computing for help with the website’s server backbone. We thank Arne Sahm for critical reading of the manuscript.

## Author Contributions

M.F. conceived the study. M.F. and S.H. supervised the work. M.F. and K.R. curated and analyzed the data. R.S., K.R., M.F., and S.H. designed the website. R.S. created the website. M.F., J.A.D., and S.H. interpreted the data. M.F., with the help of S.H. and J.A.D., wrote the manuscript. All authors read and approved the manuscript.

